# CRISPR-Cas9 gene editing in the agriculturally beneficial entomopathogenic nematode *Steinernema feltiae*

**DOI:** 10.1101/2025.07.18.665633

**Authors:** Sally W. Ireri, Mengyi Cao

## Abstract

The entomopathogenic nematodes (EPN) of the genus *Steinernema* serves as a valuable experimental model for studying microbial symbiosis and is economically important as organic pest control agent in agriculture. Although most *Steinernema* species are dioecious (male-female), consistent genetic manipulation has thus far only been demonstrated in the hermaphroditic species *Steinernema hermaphroditum*. In this study, we adapted a CRISPR-Cas9–based gene editing approach to *Steinernema feltiae*, a dioecious species widely used in agricultural applications. Using gonadal microinjection, we targeted the conserved gene *unc-22* in *S. feltiae* and generated stable mutant lines with large on-target deletions. Mating tests revealed that *Sf-unc-22* is X-linked and exhibits a conditionally dominant twitching phenotype. Additionally, *Sf-unc-22* mutants display distinct body morphology compared to wild-type nematodes. Homozygous mutant lines can be reliably maintained through cryopreservation. Altogether, our work provides a proof of concept that genetic tools developed in *S. hermaphroditum* can be effectively adapted to other agriculturally relevant and dioecious *Steinernema* species— broadening the scope of molecular genetic research in microbial ecology and enhancing their potential applications in agriculture.

## Introduction

Steinernematids are soil-dwelling entomopathogenic (EPN, insect-parasitic) nematodes that form a symbiotic relationship with gram-negative bacteria from the genus *Xenorhabdus* ^1,2^. The nematodes are typically obligate insect parasites throughout most of their lifecycle, except in the non-feeding infective juvenile (IJ) stage where they leave the insect and persist in the soil ^3^. The IJs are associated with species-specific symbiotic *Xenorhabdus* bacteria in an intestinal pocket, known as the receptacle ^4^. Once a new insect host is found, the IJs invade the insect through natural orifices, releasing their symbiotic bacteria which cause the death of the insect *via* septicaemia ^5,6^. Within the insect host, the nematodes exit the arrested-development IJ stage and complete their life cycle while feeding on their symbiont for several generations ^7,8^. Eventually, depletion of nutrients and overcrowding triggers the formation of IJs which reassociate with their symbiont and disperse to seek a new insect host ^3,9^.

The *Steinernema feltiae - Xenorhabdus bovienii* symbiotic pair has been applied in home gardens (small-scale) and commercial agriculture (large-scale) since the 1970s, and proven to be a useful tool for integrated pest management (IPM) ^10^. *S. feltiae* represents a particularly potent and attractive EPN due to the relatively low concentrations of IJs necessary for effective pest control compared to other EPNs ^11^, as well as relatively high potency against insect pests ^6^. *S. feltiae* IJs parasitize a wide variety of insects, with successful application against soil-dwelling pests of grains, berries, vegetables, mushrooms, fruit trees, and tuber crops, among others ^5,12,13^. Typically in large-scale commercial applications, EPNs are delivered as a suspension in a liquid medium, sometimes mixed with chemical pesticides ^14^. Currently, multiple challenges limit the broader commercial uptake of EPNs as biological control agents. One such challenge is the inconsistency in efficacy and shelf life, which results in mixed outcomes following application. Additionally, persistence of EPNs in the field for multiple seasons after application is hindered by their susceptibility to UV radiation and desiccation ^13^. These challenges illustrate the urgent need to better understand the molecular mechanisms underlying the infectivity dynamics in the field ^5,15,16^ and develop molecular genetic tools that complement existing strategies to enhance EPNs tolerance to abiotic stressors ^14,17^.

While the bacterial partner, *X. bovienii,* is genetically tractable *via* multiple techniques such as conjugation and CRISPR-Cas systems ^18–21^, no genetic tool is available to study the *S. feltiae* nematode host (reviewed in ^22^), limiting the understanding of basic biology of this species and its application in agriculture. Recently, consistent CRISPR-Cas9 gene-editing technology has been proven successful in *Steinernema hermaphroditum,* a hermaphroditic species that is not currently applicable in agriculture. It is crucial to develop consistent genetical tools in an agriculturally relevant species of *Steinernema* as these nematodes are exclusively male-female and may encounter challenges that hermaphroditic species do not encounter. For instance, although temporary gene editing was observed to be successful in a male-female and agriculturally relevant species *Steinernema carpocapsae*, creating homozygous alleles and maintaining mutant lines was unsuccessful due to the difficulties in microinjection, recovery, breeding *in vitro* and *in vivo,* and cryopreservation ^23^.

In the model organism *Caenorhabditis elegans, unc-22* encodes a protein known as Twitchin that is involved in muscle contraction, and has been used as a classical genetic marker where the loss-of-function allele results in a distinctive phenotype of impaired movement, known as twitching ^24,25^. Since the twitching phenotype in heterozygous animals is conditionally dominant in the presence of nicotine, *unc-22* alleles can be used as a co-CRISPR marker to enrich other mutations that are not easily scored phenotypically ^23,26^. The *unc-22* homologues are highly conserved across nematode species ranging from the human intestinal parasite *Strongyloides stercoralis* ^27^ to the insect parasitic nematode *Steinernema hermaphroditum* ^23^.

Here we report the first case of consistent gene editing in *S. feltiae,* an EPN species that is dioecious and agriculturally relevant. We targeted an *unc-22* homologue in *S. feltiae* to establish a dominant allele as a genetic marker in this species, and as proof of concept for successful CRISPR-Cas9 gene editing. Both wild-type *S. feltiae* and the *Sf-unc-22* mutant recovered well from cryopreservation which facilitates long-term storage and genetics research. Additionally the nematodes develop optimally at room temperature which is convenient for rearing the animals in the laboratory ^28^. Since *S. feltiae* is a beneficial organism broadly applicable in the field, this technique is crucial to the study of gene function in *S. feltiae* that may prove to be important in improving its efficacy in agriculture. Altogether, with its value as an experimental system for mutualistic and parasitic symbioses, the *S. feltiae - X. bovienii* partners serve as a promising emerging genetic model.

## Materials and Methods

### Bacterial growth and nematode maintenance

*Steinernema feltiae* Florida (FL) wild-type conventional infective juveniles (IJs) ^1^ (MCN0013) were maintained through *Galleria mellonella* 5^th^ instar insects (Grubco, OH) by white-trapping ^29^. *Steinernema feltiae* wild-type and mutant lines were maintained at room temperature (approximately 22°C) on Nematode Growth Media (NGM) seeded with *X. bovienii - Sf-*FL (MCB038). Bacteria *Xenorhabdus bovineii-Sf-*Florida were grown at 30°C on Luria Bertani (LB) agar supplemented with 0.1% sodium pyruvate ^30^. A single colony was picked to start overnight culture in dark LB media and grown at 30°C with aeration. *Comammonas acquatica* (DA1877) was grown at 37°C on LB agar or liquid media with aeration and seeded on NGM plates for nematode post-injection recovery.

### Target gene sequence analysis and Cas9 ribonucleoprotein preparation

The *Sf-unc-22* homologue was determined using the BLAST algorithm in WormBase ParaSite (Version: WBPS19) and identified as gene ID contig03263.2.54 ^31–33^. Primers were used to amplify a segment of the gene that was highly conserved with the successfully edited exon of the *Sh-unc-22* homologue ^23^. The sequence of the target segment was confirmed using Oxford-Nanopore sequencing (https://plasmidsaurus.com/) before CRISPR RNAs (crRNAs) were designed using the web tool CHOPCHOP Bio (https://chopchop.cbu.uib.no/) ^34^. The top candidates, ranked by the algorithm, were aligned against the *S. feltiae* NW Washington strain genome to check for potential off-target matches. The top 4 unique crRNAs (rank number 1, 2, 7 and 9) were chosen and ordered (Integrated DNA Technologies, Inc., Coralville, IA) (Table S1). To make the Cas9 ribonucleoprotein, *C. elegans* and *S. hermaphroditum* CRISPR protocols ^23,35^ were adapted as follows: 1.5 μL of each crRNA (100 μM) is mixed with tracrRNA (100 μM) at a 1:1 ratio in 0.2 μL PCR tubes. The mix is heated to 94℃ in a thermocycler for 2 minutes then cooled to room temperature. To make the injection mix, 1.16 μL of 1M KCl was added to the duplex made above, followed by 4 μL of Cas9 protein (Integrated DNA Technologies, Inc., Coralville, IA). The injection mix was incubated at room temperature for 5 minutes then mixed three times by tapping and spun down, with a final spin down of 1 min. The injection mix was placed on ice and microinjection followed immediately.

### Gonadal microinjection of *S. feltiae*

*S. feltiae* nematodes were grown on NGM plates on a patch of *X. bovineii*. A young adult female nematode was picked and rinsed in M9 buffer before being placed in a drop of halocarbon oil 700 (Sigma-Aldrich, Inc., St. Louis, MO) on a 2% agarose pad. Microinjection was carried out targeting each gonad if possible, using quartz needles to deliver the injection mix *via* the CLEAN function on the FemtoJet 4x (Calibre Scientific, Los Angeles, CA) at approximately 99 psi injection pressure. The injected nematode was immediately recovered using a drop of M9 on the agarose pad, rinsed to remove the oil, and placed on an individual NGM plate seeded with *C. aquatica.* The injected nematodes (P0) were monitored for survival of microinjection then 3-5 males were added to the plate to allow for continuous mating. The P0 nematodes were allowed to recover on individual *C. aquatica* plates at room temperature. F1 progeny were screened with 2% nicotine to identify the heterozygous twitchers.

### Genotyping and Sanger sequencing to confirm on-target genome editing

Pam2 and Pam7 were combined into one genotyping reaction using primers Sf_unc22_For and Sf_unc22_cr7_Rev, while Pam1 and Pam9 target sites were similarly then genotyped using primers Sf_unc22_cr1_For and Sf_unc22_cr9_Rev. For a full list of primers used in this study see Table S2.

### Confocal microscopy and image analysis

Wild-type and *unc-22* mutant nematodes at J4 stage were imaged by confocal microscopy LSM900 (Zeiss, Jena, Germany). Images were processed by ZenPro software and quantified in Fiji ^36^. Specifically, the length (head to tail) and width (vertical to vulva) of *Sf-unc-22* and wild-type nematodes were measured by segmented lines. The statistical analysis of paired t-tests was performed in Prism 10 (GraphPad Software, MA).

### Determining linkage group of *Sf-unc22* homologue

Briefly, six *Sf-unc-22* virgin females at J4 stage were placed on a lawn of symbiotic bacteria *X. bovienii* on an NGM plate. Twice the number of wild-type *S. feltiae* males were added to the plate and the nematodes were allowed to mate for three days at room temperature. Productive females with visible progeny (usually shown as bagging) were separated on individual NGM plates seeded with *X. bovienii*, and the progeny were allowed to develop to J3 or J4 stage before being screened for twitching in the presence and absence of 2% nicotine. Animals twitching in the absence of nicotine are scored as haploid or homozygous; animals that are twitching in the presence of nicotine are scored as heterozygous *unc-22/*+. Animals that do not twitch in nicotine are scored as +/+.

### Cryopreservation

Cryopreservation of *S. feltiae* lines was performed using trehalose-DMSO freezing solution, a method adapted from *S. hermaphroditum* and *C. elegans* ^37,38^. To test the efficiency of cryopreservation of wild-type, *mc0014,* and *mc0015* mutant strains, three replicates of frozen stocks of each strain (each containing 1 mL of nematode samples) were stored at -80°C. One month after storage, each frozen stock was thawed and recovered on an individual NGM agar plate at room temperature (22°C). A reproductive thaw is scored when nematodes recovered on the NGM plates produced progeny.

## Results

### Gonadal microinjection delivers CRISPR-Cas9 ribonucleoprotein in *S. feltiae* and successfully modifies *Sf-unc-22* homologue

We identified an *unc-22* homologue in *S. feltiae* by aligning *C. elegans unc-22* against the *S. feltiae* genome using the BLAST algorithm ^32,33^. This homologue showed 89.6% identity to the amino acid sequences of the *S. hermaphroditum unc-22* homologue (*Sh-unc-22*) (Fig. 1A). Exon 40 of *Sf-unc-22* with high homology to an exon that was previously mutated in *S. hermaphroditum unc-22* by CRISPR-Cas9 was identified and used as target site in *S. feltiae* (Fig. 2A). Four crRNAs (Table S1) were incorporated into the injection mix to increase the chances of successful gene editing ^23^.

**Figure 1:**
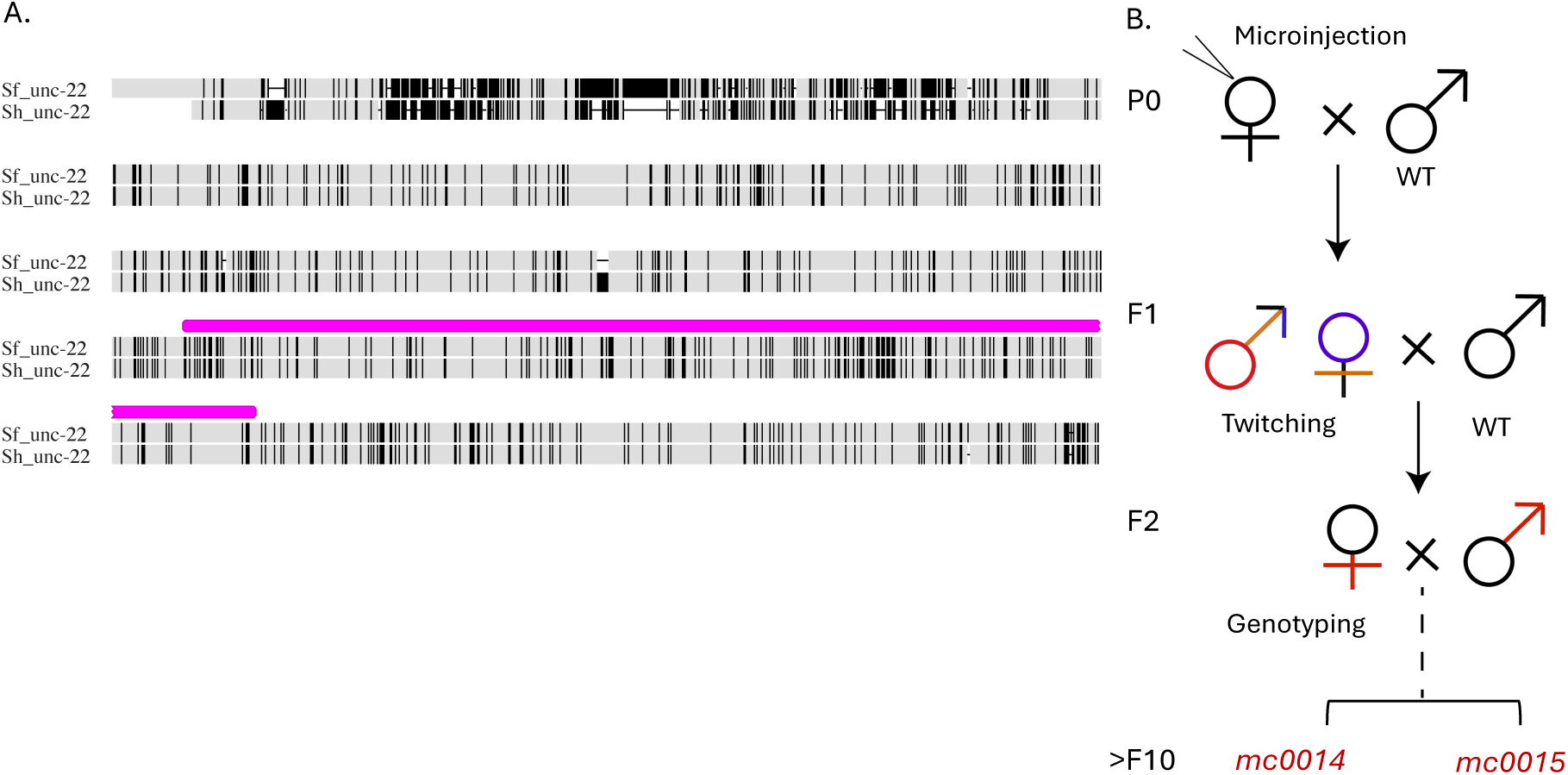
Microinjection of *S. feltiae* targeting the *sf-unc-22* homologue. (A): Protein alignment of the Sf-UNC-22 and Sh-UNC-22 homologues with the Cas9 target region indicated by a pink bar on top of the alignment. The amino acid sequences were pairwise aligned in Geneious (Geneious Prime® 2025.1.3) using the Geneious Alignment algorithm for a global alignment with free end gaps and Blosum62 cost matrix. All disagreements to the consensus sequence are highlighted in black, including ambiguous disagreements**. (**B): Flowchart showing the mating process involved in creating the stable transgenic lines MCN0014 and MCN0015. The P0 female was allowed to mate with wildtype (WT) males, resulting in genetically mosaic F1 progeny with a variety of alleles due to repair of the Cas9-induced double-stranded breaks. Twitching F1 females were backcrossed with WT males. The progeny, F2, were self-mated for several generations resulting in two stable alleles, *mc0014* and *mc0015*.

**Figure 2:**
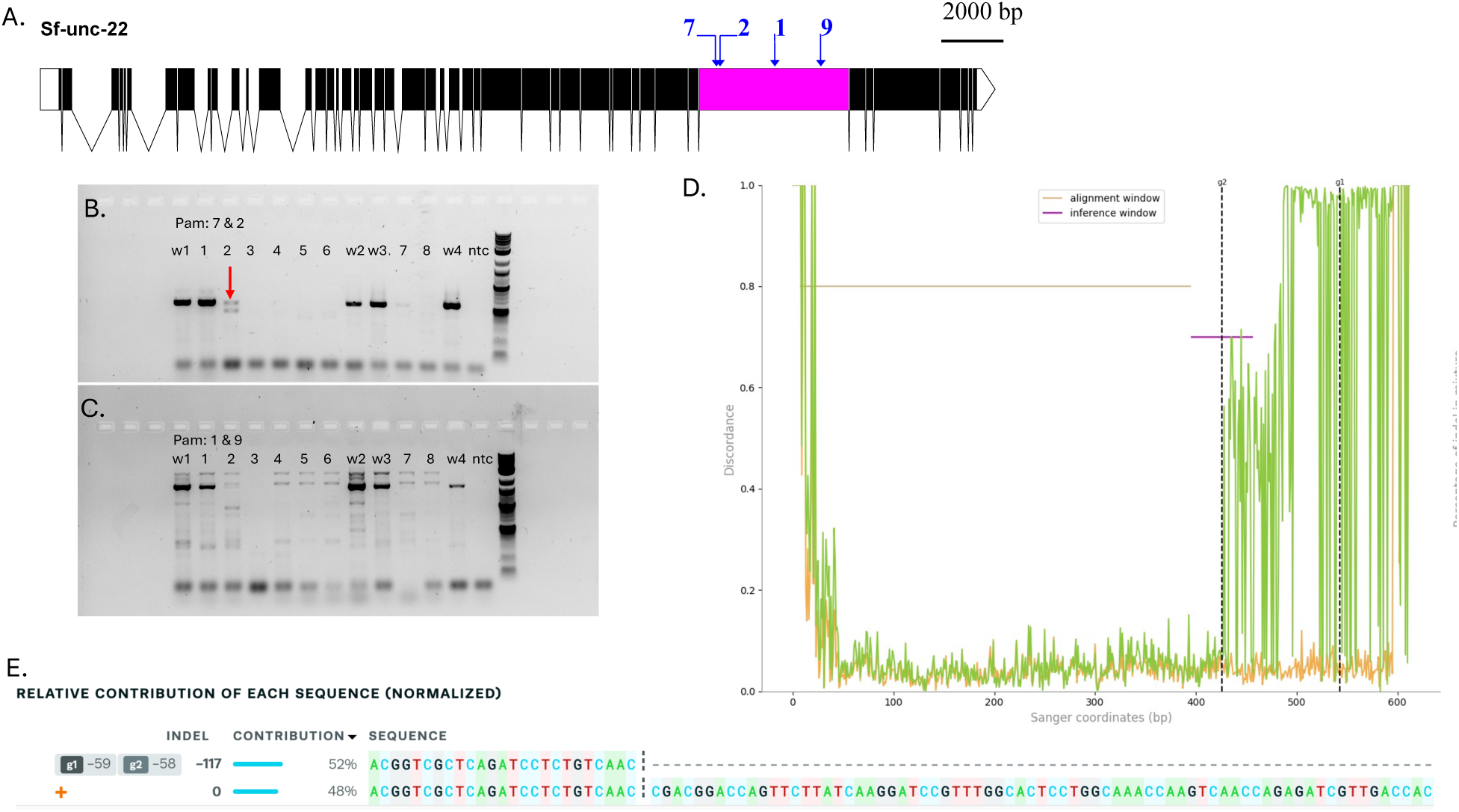
Analysis of F2 twitching nematodes. (A): Gene structure of *Sf-unc-22* showing Pam sites in order: Pam7, Pam 2, Pam1, Pam9. (B) and (C): Genotyping PCR of F2 individuals. (B) shows the Pam7 and Pam2 genotyping PCRs, while (C) shows the Pam1 and Pam9 genotyping PCRs. 1.2% agarose, NEB 1kb+ ladder. W1, w2, w3, w4 – wildtype, ntc – non-template control. (See supplementary table S3 for lineage information of each F2 sample). (D): Trace analysis of chromatograms from Sanger sequencing of amplicons of the Pam7 and Pam2 target sites. Analysis was carried out using the Inference of CRISPR Edits software (ICE). The left panel shows good alignment between the wildtype (WT) chromatogram (in orange), and the chromatogram of sample 2 (in green) in figure 2B indicated by a red arrow. The dotted vertical lines labelled g1(Pam7) and g2 (Pam2) indicate the putative cut sites by Cas9 in the gRNA recognition site. The chromatograms of WT and sample 2 vary greatly after the g2 dotted line, indicating a divergence in the sequence from WT. Editing efficiency at the Pam2 target site is 52% with an R^2^ of 99%. (E): ICE analysis of Sanger sequencing of PCR amplicon shown by the red arrow in Fig.2D indicating the presence of two alleles: wildtype allele indicated by a “+”, and a 117bp deletion allele target site represented by the horizontal dotted line when sample 2 sequence is aligned against WT sequence.

The CRISPR-Cas9 ribonucleoprotein (RNP complex) was delivered to the female nematode gonadal syncytium via microinjection using a similar method described in ^23^. We injected pre-mated female *S. feltiae* adults with fertilized embryos present in the gonads since they survived better post-injection than younger adults. Additionally, we used quartz needles rather than borosilicate needles which better penetrates the relatively thick cuticle of adult females. All 12 injected nematodes (P0) showed immediate signs of recovery confirmed by pharyngeal pumping within approximately 30 minutes to an hour after injection, when recovering on individual lawns of *C. aquatica* (Table 1). Wild-type *S. feltiae* males were added to the recovery plates to allow for continuous mating overnight. All 12 injected females survived overnight (Fig. 1B) and 10 produced F1 progeny (Table 1).

**Table 1:**
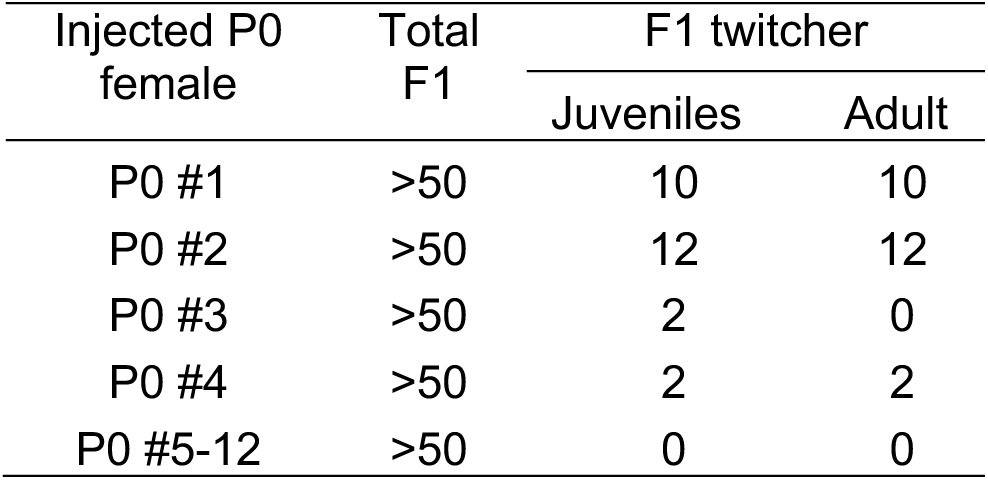
*S. feltiae* P0 females with twitching progeny.

The F1 nematodes were allowed to develop, tested and scored in 2% nicotine to identify conditionally dominant alleles in heterozygous animals (Table 1). Twitching female F1s were isolated on separate plates on a lawn of symbiont *X. bovienii*. Four P0 nematodes were found to have produced progeny that twitched in 2% nicotine, giving an editing frequency of one twitcher out of three injected female *S. feltiae* (33% frequency of success) (Table 1). Since many F1 twitching nematodes, especially the males, did not survive to adulthood, we backcrossed individual twitching F1 females with wild-type males and scored their progeny (F2) in 2% nicotine (Fig. 1B). Four out of the five F1 females produced progeny, with two producing twitching F2 nematodes (Table 2). Some F2 twitchers (see Table S3 for lineage information) were isolated for genotyping in case they cannot be propagated further (Fig. 2B and 2C). We detected the presence of indels in F2 twitching progeny by Sanger sequencing and analyzed the sequence chromatograms using the ICE program ^39^ (Table 3; Fig 2D, 2E).

**Table 2:** F1 twitcher breeding test.

| F1 twitching female | Parental P0 | F2 progeny | F2 twitchers |
| --- | --- | --- | --- |
| F1-1 | P0 #1 | Yes | No |
| F1-2 | P0 #2 | Yes | 12 |
| F1-3 | P0 #2 | Yes | No |
| F1-4 | P0 #4 | Yes | 2 |
| F1-5 | P0 #4 | No | N/A |

**Table 3:** Indels in *S. feltiae* F2 uncovered by Sanger sequencing.

| Label on gel | Parental Line | Sex | Generation | Mutations at pam7&2 | Mutations at pam1&9 |
| --- | --- | --- | --- | --- | --- |
| 1 | P0-1 | ♀ | F2 | - | - |
| 2 | P0-2 | ♂ | F2 | 117 bp del | - |

**Table 4:** Efficiency of recovery from cryopreservation of *S. feltiae*.

| Strain | Vials tested | Productive thaw |
| --- | --- | --- |
| Wild-type | 3 | 3/3 |
| *mc0014* | 3 | 3/3 |
| *mc0015* | 3 | 2/3 |

One F1 line (F1-2) had 12 severe twitchers and was allowed to inbreed among siblings. After more than 10 generations of inbreeding, two stable lines emerged (MCN0014, MCN0015) which have a consistently severe twitching phenotype without nicotine, suggesting they may be homozygous recessive alleles (Fig 3). Amplification of the target region of the *Sf-unc-22* homologue followed by sequencing confirmed the homozygosity and revealed the presence of large deletions of more than 3,300 base pairs beginning 4 bases upstream of the Pam 9 target site (Fig 3A, 3B), a case of on-target editing.

**Figure 3:**
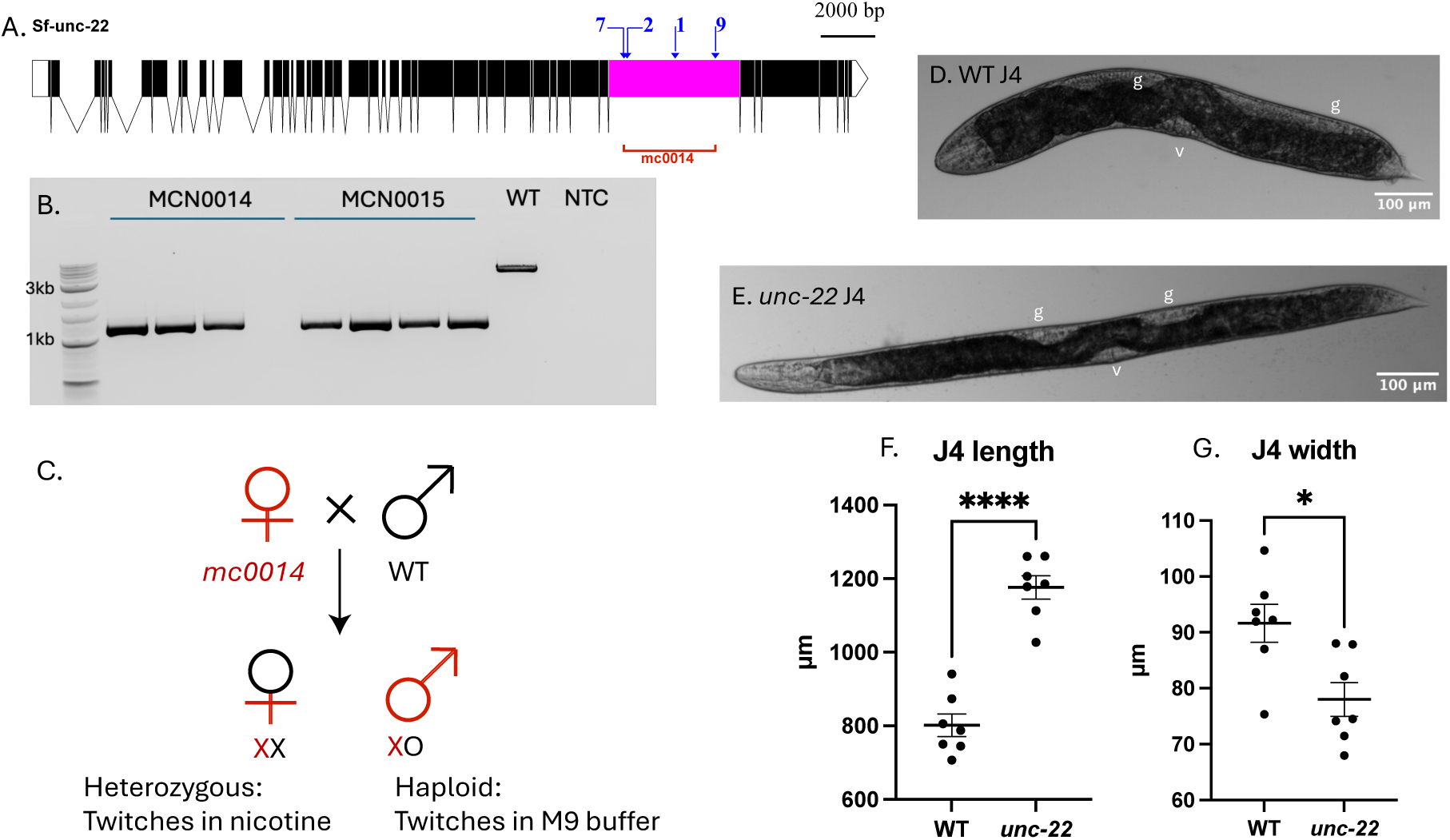
Genotyping and phenotyping of *Sf-unc-22* mutant lines. (A): Illustration of the large deletion present in one of the stable lines obtained from the CRISPR-Cas9 knockout experiments (MCN0014). The deletion is marked by a red line identified by the allele number mc0014. (B): Gel electrophoresis showing the results of amplification of the 4500 bp target region of the *sf-unc-22* homologue. A 1% agarose gel was run at 120V for 45 minutes. The wild-type (WT) amplicon serves as a positive control, while the non-template control (NTC) used molecular grade water as a negative control. Both MCN0014 and MCN0015 exhibit a homozygous deletion of 3,370 bp, and 3,371 bp respectively. This is evidenced by the absence of a WT band in the PCR amplicon and confirmed by sequencing. (C): To test whether the *Sf-unc-22* homologue is X-linked, 6 pre-mated early J4 females from the strain MCN00014 were placed with 12 wildtype males and allowed to mate for several days at room temperature. The *Sf-unc-22* females were transferred to individual NGM plates seeded with symbiont once the gonads were full of eggs. Progeny were separated by sex and tested first in M9, then in 2% nicotine. Males twitched in both M9 and nicotine, suggesting that the gene is X-linked. Females twitched only in nicotine, indicating a conditional dominant allele in the heterozygous state. (D): and (E). portray an *Sf-WT* and an *Sf-unc-22* nematode respectively. The S*f-unc-22* J4 female is relatively longer and thinner than the WT *S. feltiae* when treated with levamisole for confocal microscopy. (F) and (G): analysis of length (F) and width (G) of seven individual wild-type compared to seven *Sf-unc-22* nematodes in microns.

To further characterize the morphology of *unc-22* mutants, we imaged both *Sf-unc-22* mutant and wild-type J4 females by confocal microscopy. Image analysis showed that the *Sf-unc-22* nematodes have significantly greater body length and reduced body width when compared to wild-type animals (Fig 3D, 3E, 3F, and 3G). Since the severity of the twitching phenotype may not always be obvious depending on the allele, these morphological signatures associated with body metrics could be used as additional phenotypic markers to screen and identify *unc-22* mutants (Fig 3D and Fig 3E).

### *Sf-unc-22* homologue is X-linked in *S. feltiae*

In *C. elegans,* the *unc-22* gene is linked to Chromosome IV, while based on the genomic analysis of *S. hermaphroditum* ^23^ the *unc-22* homologue is found on the X chromosome. The *S. feltiae* genome is incomplete and chromosomal information is currently unavailable ^31^. In order to determine if the *Sf-unc-22* homologue is X-linked as in *S. hermaphroditum*, we set up a mating test using *Sf-unc-22* virgin females (XX) and wild-type males (XO) (Fig 3C). We observed F1 male progeny twitch autonomously in M9 buffer (without nicotine treatment), confirming they are haploid (X*^unc-22^*O). Meanwhile, F1 female progeny twitch in 2% nicotine exclusively, showing they are heterozygous, carrying the *unc-22* conditionally dominant allele (X*^unc-22^*X). These results are consistent with our hypothesis that *unc-22* is linked to the X chromosome in *S. feltiae,* as it is in other *Steinernema* species ^23,40^.

## Discussion

In this research, we show the first successful case of consistent CRISPR-Cas9 mediated gene editing in a dioecious species of entomopathogenic nematode. We demonstrate that genome editing technology developed in a hermaphroditic model organism of Steinernematids provides a launch-pad for optimization of CRISPR-Cas9 mediated editing in other *Steinernema* species, most of which are strictly dioecious. We also show that *unc-22* is highly conserved between *S. feltiae* and *S. hermaphroditum,* both by amino acid sequence alignments (Fig. 1A) and by the mutant twitching phenotypes. The conservation of this gene and its mutant phenotype across *Steinernema* species makes the *unc-22* homologue a good marker for genetic perturbation studies in this family of nematodes and provides support for the use of *unc-22* as a co-CRISPR marker to aid in the development of transgene insertion protocols.

The *S. feltiae unc-22* mutants created by CRISPR-Cas9 genome editing can be further crossed and inbred to produce homozygous mutant lines and can be cryopreserved. The availability of breeding and cryopreservation will greatly facilitate further development of genetic tools in *S. feltiae.* Interestingly, preliminary results show that the *S. feltiae unc-22* strains obtained from this study are unable to be recovered from natural propagation through insects, suggesting a potential defect in infection, and/or reproduction in the insect host. This is different from *S. hermaphroditum unc-22* mutants which have been shown to successfully infect *Galleria mellonella*, reproduce in the insect, and subsequently be recovered by White trapping ^23^.

The ability to precisely knock out genes in *S. feltiae* will advance research in genetics, parasitology, and microbial symbiosis by enabling detailed functional studies of nematode genes. Genetic tractability of both partners in the *S. feltiae*–*X. bovienii* symbiosis is essential for dissecting the molecular pathways that underpin their mutualistic relationship. Thus, the capacity to genetically manipulate *S. feltiae* opens exciting opportunities to investigate the complex biology of the tripartite interaction among the nematode, its bacterial symbiont, and the insect host.

CRISPR-Cas9–mediated genome engineering offers a powerful approach to enhance the efficacy of *S. feltiae* as a biological control agent in agricultural settings. To date, efforts to improve field performance of entomopathogenic nematodes (EPNs) have primarily relied on traditional selective breeding methods ^41^ ^42^. While these approaches have yielded moderate success— particularly when combined with innovations in application methods such as deploying infected cadavers as delivery vehicles ^14^—CRISPR-Cas9 enables precise genetic edits that can target specific traits beneficial for field performance. Furthermore, deeper insights into key aspects of nematode biology, including foraging behavior ^43^ and dispersal cues in infective juveniles ^44^ will facilitate more targeted and effective pest management strategies. To fully realize the potential of CRISPR-Cas9 in *S. feltiae*, additional work is needed to establish robust protocols for the precise insertion of genetic cargo into its genome.

## Acknowledgements

We thank Heidi Goodrich-Blair for sharing *S. feltiae* and *X. bovienii* strains used in this research. We thank Margaret McFall-Ngai for sharing the confocal microscope and the needle puller and supplies for microinjection. We thank Stephanie Hampton and John Mulchaey for allocating funds to support the microinjection equipment. We thank Zoila Jurado Quiroga for help assembling the microinjection equipment.

## Funding

This research is supported by Carnegie Institution for Science.

## Author contributions

SWI and MC initiated and conceptualized the project, developed methods, performed experiments, and managed the project. SWI designed crRNA, microinjection, and performed sequencing data analysis independently. SWI wrote the original draft. SWI and MC edited the manuscript.

## Supplemental materials

**Table S1:**
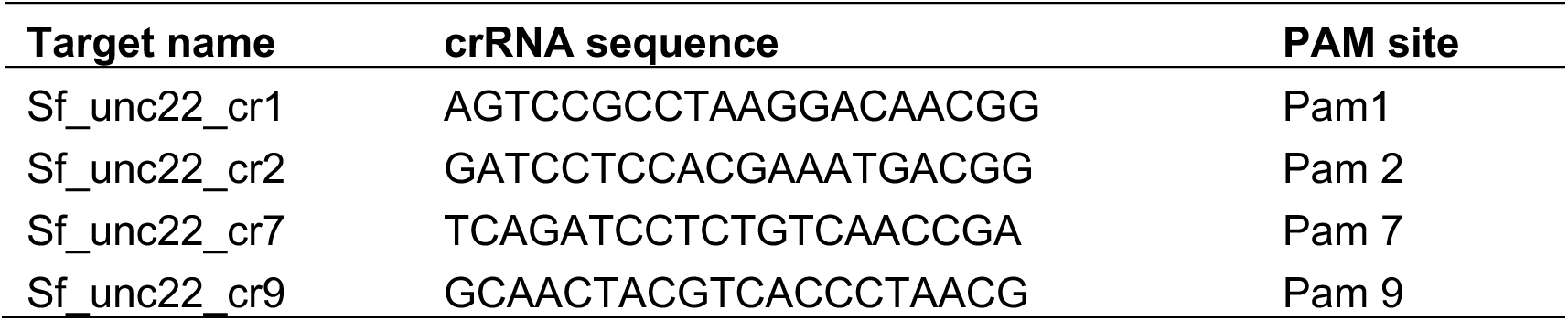
A list of guide RNAs used in this research.

**Table S2:**
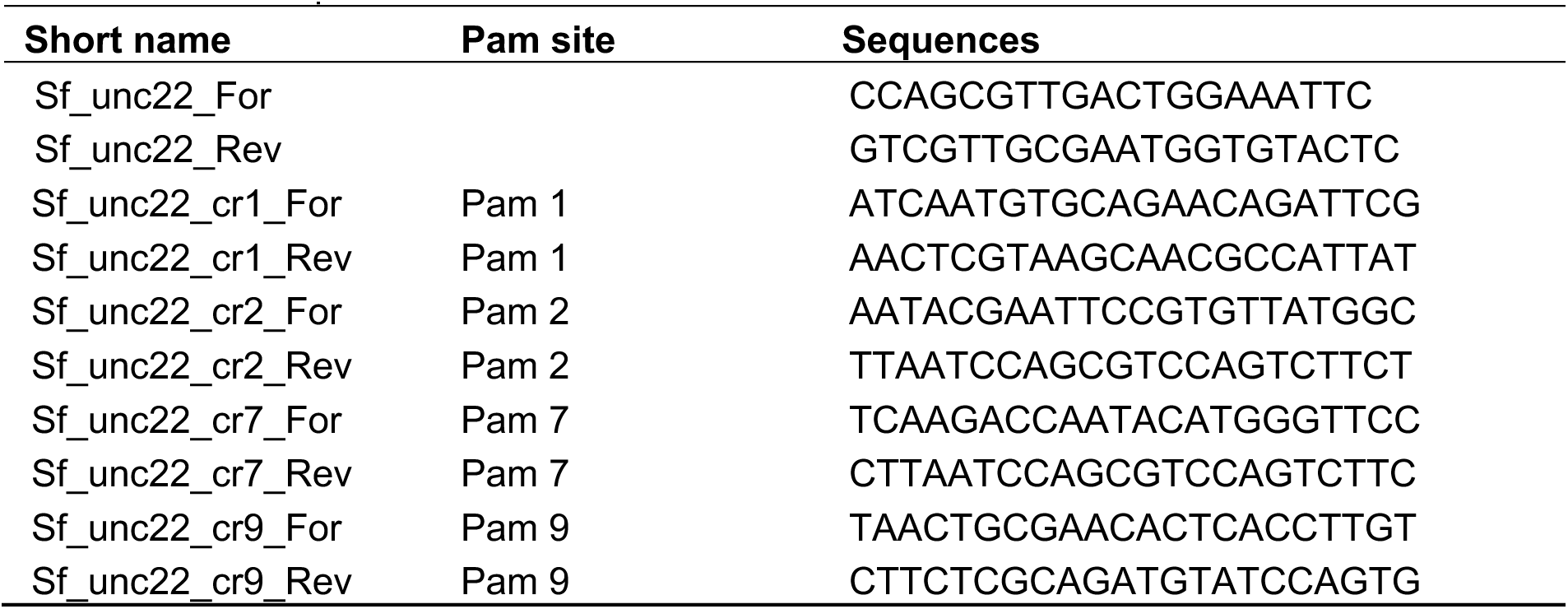
A list of primers used in this research.

**Table S3:** Lineage information of F2 on gel shown in Fig. 2B and 2C

| Label on gel | Parental Line | Sex | Generation |
| --- | --- | --- | --- |
| W1 | WT | ♀ | - |
| 1 | P0-1 | ♀ | F2 |
| 2 | P0-2 | ♂ | F2 |
| 3 | P0-2 | ♂ | F2 |
| 4 | P0-2 | ♂ | F2 |
| 5 | P0-2 | juv | F2 |
| 6 | P0-2 | ♂ | F2 |
| W2 | WT | ♂ | - |
| W3 | WT | ♀ | - |
| 7 | P0-2 | 2 juv | F2 |
| 8 | P0-2 | 2 juv | F2 |
| W4 | WT | - |  |

## Notes

### Competing Interest Statement

The authors have declared no competing interest.

